# EEG-Informed fMRI Analysis Reveals Neurovascular Coupling in Motor Execution and Imagery

**DOI:** 10.1101/2025.01.13.632624

**Authors:** Elena Bondi, Yidan Ding, Yisha Zhang, Eleonora Maggioni, Bin He

## Abstract

The complementary strengths of electroencephalography (EEG) and functional magnetic resonance imaging (fMRI) have driven extensive research into integrating these two noninvasive modalities to better understand the neural mechanisms underlying cognitive, sensory, and motor functions. However, the precise neural patterns associated with motor functions, especially imagined movements, remain unclear. Specifically, the correlations between electrophysiological responses and hemodynamic activations during executed and imagined movements have not been fully elucidated at a whole-brain level. In this study, we employed an EEG-informed fMRI approach on concurrent EEG-fMRI data to map hemodynamic changes associated with dynamic EEG temporal features during motor-related brain activities. We localized and differentiated the hemodynamic activations corresponding to continuous EEG temporal dynamics across multiple motor execution and imagery tasks. Validation against conventional block fMRI analysis demonstrated high precision in identifying regions specific to motor activities, underscoring the accuracy of the EEG-driven model. Beyond the expected sensorimotor activations, the integrated analysis revealed supplementary coactivated regions showing significant negative covariation between blood oxygenation level-dependent (BOLD) activities and sensorimotor EEG alpha power, including the cerebellum and insular cortex. These findings confirmed both the colocalization of EEG and fMRI activities in sensorimotor regions and a negative covariation between EEG alpha band power and BOLD changes. Moreover, the results provide novel insights into neurovascular coupling during motor execution and imagery on a brain-wide scale, advancing our understanding of the neural dynamics underlying motor functions.

**Highlights:**

- Movement-related EEG alpha activity negatively covaries with BOLD responses
- EEG-informed fMRI maps the neurovascular correlates across the entire brain
- A unified model disentangles multiple motor-related brain activations
- EEG-informed fMRI results highly overlap with conventional block fMRI results

## 1. Introduction

Electroencephalography (EEG) and functional magnetic resonance imaging (fMRI) offer valuable insights into a broad spectrum of cognitive, sensory, and motor functions individually with high temporal resolution for EEG and high spatial resolution for fMRI. These complementary attributes have driven extensive research into electrophysiological-hemodynamic coupling across various brain activities, aimed at achieving high spatio-temporal resolution for human brain mapping (Case et al., 2016; He & Liu, 2008; Huster et al., 2012; Liu & He, 2008; Liu et al., 2009; Liu et al., 2010; Maggioni et al., 2016 Ritter & Villringer, 2006).

EEG measures scalp-level electrical potential fluctuations arising from synchronized neural activity, offering millisecond-scale temporal resolution critical for investigating rapid neural dynamics. Oscillatory EEG activity across specific frequency bands—delta, theta, alpha, beta, and gamma—has been linked to distinct cognitive, sensory, and motor processes. The suppression of alpha and beta rhythms over the sensorimotor region is found to be closely associated with motor actions and motor imagery (MI) (Pfurtscheller & Lopes da Silva, 1999). These neurophysiological markers underpin the development of MI-based Brain-Computer Interface (BCI) systems, which enable control of external devices by translating imagined motor tasks into control commands, circumventing peripheral neuromuscular pathways. EEG-based BCIs are portable, cost-effective, and enable real-time monitoring of brain activity, making them a key component of non-invasive neurotechnology (Edelman et al., 2024; He et al., 2020; McFarland & Wolpaw, 2017; Wolpaw et al., 2002). They have demonstrated effectiveness across a range of users, from those with motor impairments to able-bodied individuals, showcasing their versatility and potential for clinical and home use (Ang et al., 2015; Edelman et al., 2019; Forenzo et al., 2024; LaFleur et al., 2013; Meng et al., 2016; Tonin et al., 2022).

While EEG enjoys high temporal resolution, it has limited spatial resolution and suffers from a low signal-to-noise ratio due to the volume conduction effect, which disperses electrical signals as they propagate to the scalp (He et al., 2018). In contrast, b lood oxygenation level-dependent (BOLD) fMRI offers high spatial resolution, enabling precise spatial mapping of brain activities through hemodynamic responses related to neuronal activation. The somatotopic mapping of the primary motor cortex during both executed and imagined movements has been well studied using fMRI (Ehrsson et al., 2003; Meier et al., 2008; Szameitat et al., 2007), significantly enriching our understanding of motor processes and their neural underpinnings. However, as fMRI reflects fluctuations in blood oxygenation tied to neural mass activity and corresponding metabolic demands, it serves as an indirect indicator of neural activity rather than directly capturing neurophysiological dynamics. This distinction leaves critical gaps in our comprehension of the neural correlates bridging electrophysiological and hemodynamic responses within the motor domain, particularly in the context of movements or MI involving different body parts. A multimodal approach integrating EEG and fMRI is imperative for resolving these gaps, facilitating a more comprehensive understanding of hemodynamic responses and motor-related brain mechanisms (Logothetis, 2008; Warbrick, 2022).

Yuan et al. (2010) identified a spatial correspondence and negative covariation between task-related EEG alpha/beta-band activity and BOLD responses in the sensorimotor cortex during MI and actual movement. However, in their study, EEG and fMRI data were collected in separate sessions, which may introduce variability in brain activity due to shifts in cognitive and physiological states.

Over the past decades, technological advances have enabled simultaneous EEG-fMRI recordings, with significant analytical progress allowing for their application across various research fields (Abreu et al., 2018; Huster et al., 2012; Maggioni et al., 2023; Ritter & Villringer, 2006; Warbrick, 2022 ). The concurrent recording enables the investigation of the real-time interactions between synchronized electrical and hemodynamic signals while avoiding potential inter-session variability. Enhanced signal processing techniques have also been developed to effectively reduce MR-induced artifacts in EEG data, thereby benefiting integrated analyses. Using simultaneous EEG-fMRI signals, Zich et al. (2015) demonstrated an inverse correlation between the amplitude of contralateral sensorimotor EEG rhythms and fMRI activation in corresponding sensorimotor regions during motor tasks based on a correlation analysis of EEG power changes and concomitant BOLD signals. Similarly, Formaggio et al. (2010) found a negative correlation between EEG alpha and beta power change and the BOLD changes in the contralateral motor cortex and a positive correlation on the ipsilateral side during motor imagery. However, correlation-based analyses fail to capture the temporal dynamics of brain responses and often lack whole-brain coverage for identifying neural sources, highlighting the need for more advanced analytical approaches.

On the other hand, EEG-informed fMRI analysis offers a more integrated approach by leveraging EEG temporal patterns to inform fMRI data analysis. By guiding fMRI analysis with EEG-derived activity patterns, this method enables mapping of EEG-related activity throughout the brain at a finer spatial-temporal resolution (Abreu et al., 2018; Murta et al., 2015). Sclocco et al. (2014) employed EEG-informed fMRI during a right-hand movement task to capture the relationship between EEG rhythm dynamics (used as independent variables) and BOLD responses (used as dependent variables). Despite prior investigation on electrophysiological-hemodynamic integration in the context of motor activities, it remains unclear whether neural activations across diverse cortical regions can be effectively captured and differentiated within a unified EEG-informed fMRI analysis framework. The aforementioned studies have predominantly focused on single-task conditions, limiting the demonstration of the generalizability of such joint analyses in disentangling multiple brain activities underlying separate tasks. Furthermore, while previous research has mainly focused on the neurovascular coupling during executed movements, the neural dynamics underlying imagined movements are less well characterized than those of actual motor execution.

To address these gaps in EEG-fMRI research on motor functions, we aim to investigate neurovascular coupling across multiple task conditions, including motor execution (ME) and MI for both the right and left hands, with simultaneous EEG and fMRI recordings. Our study began with separate analyses of each modality to replicate established neuronal and hemodynamic responses elicited by these tasks. We then employed an EEG-informed fMRI approach to map hemodynamic changes associated with dynamic EEG temporal features. Using a data-driven approach across multiple ME and MI task conditions, we leveraged the continuous temporal dynamics in EEG activity to identify the corresponding hemodynamic correlates. We hypothesize that this integrated approach can identify both colocalized neural activations and coactivated regions that covary with EEG alpha power across various motor tasks involving ME and MI, offering deeper insights into the interplay between electrophysiological and hemodynamic response.

## 2. Methods

### 2.1 Subjects

Thirty-seven healthy, able-bodied participants were recruited for this study, each providing written informed consent under a protocol approved by the Institutional Review Board of Carnegie Mellon University (protocol number: STUDY2017_00000548). All participants completed three to five EEG training sessions to complete a discrete 1-dimensional (1D) cursor control task as an initial learning phase. During this period, two participants withdrew due to scheduling conflicts, and sixteen were excluded based on unsatisfactory task performance (percent valid correct < 70%). Nineteen participants progressed to a single session in which EEG and fMRI were recorded simultaneously. Data from two additional participants were excluded from the analysis due to excessive head motion and not completing the entire protocol, respectively. The concurrent EEG-fMRI data from the remaining seventeen participants (five male, twelve female; aged 23.94 ± 3.33 years; fifteen right-handed) were included in the final analyses.

### 2.2 Experimental Protocol

The experimental protocol was designed using PsychoPy® software (Peirce et al., 2019) and consisted of alternating task blocks, each lasting 22 seconds, and resting-state blocks, each lasting 16 seconds, as shown in Fig. 1. Task blocks involved either ME or MI conditions. Each task block began with a 2-second text instruction indicating the upcoming task, followed by four task trials, each lasting 3 seconds and interleaved with 2-second inter-trial intervals. Task conditions included three ME tasks (left hand, right hand, right foot) and three corresponding MI, with ME blocks always preceding MI blocks for the same movement. No EEG activity was observed for right foot movement or imagery. Consequently, these two task conditions were excluded from further analysis. Tasks within each block were kept the same, while block sequences were randomized and balanced across sessions. During task blocks, participants were instructed to repeatedly clench their left hand, right hand, or right toes at a rate of 2 Hz, or to imagine clenching these body parts at the same pace, guided by a visual cue that flashed at the target speed. All participants lay supine with their arms relaxed, and their heads were stabilized with sponge paddings on both sides. They were instructed to keep their heads as still as possible and minimize unnecessary movements throughout the experiment.

**Figure 1.**
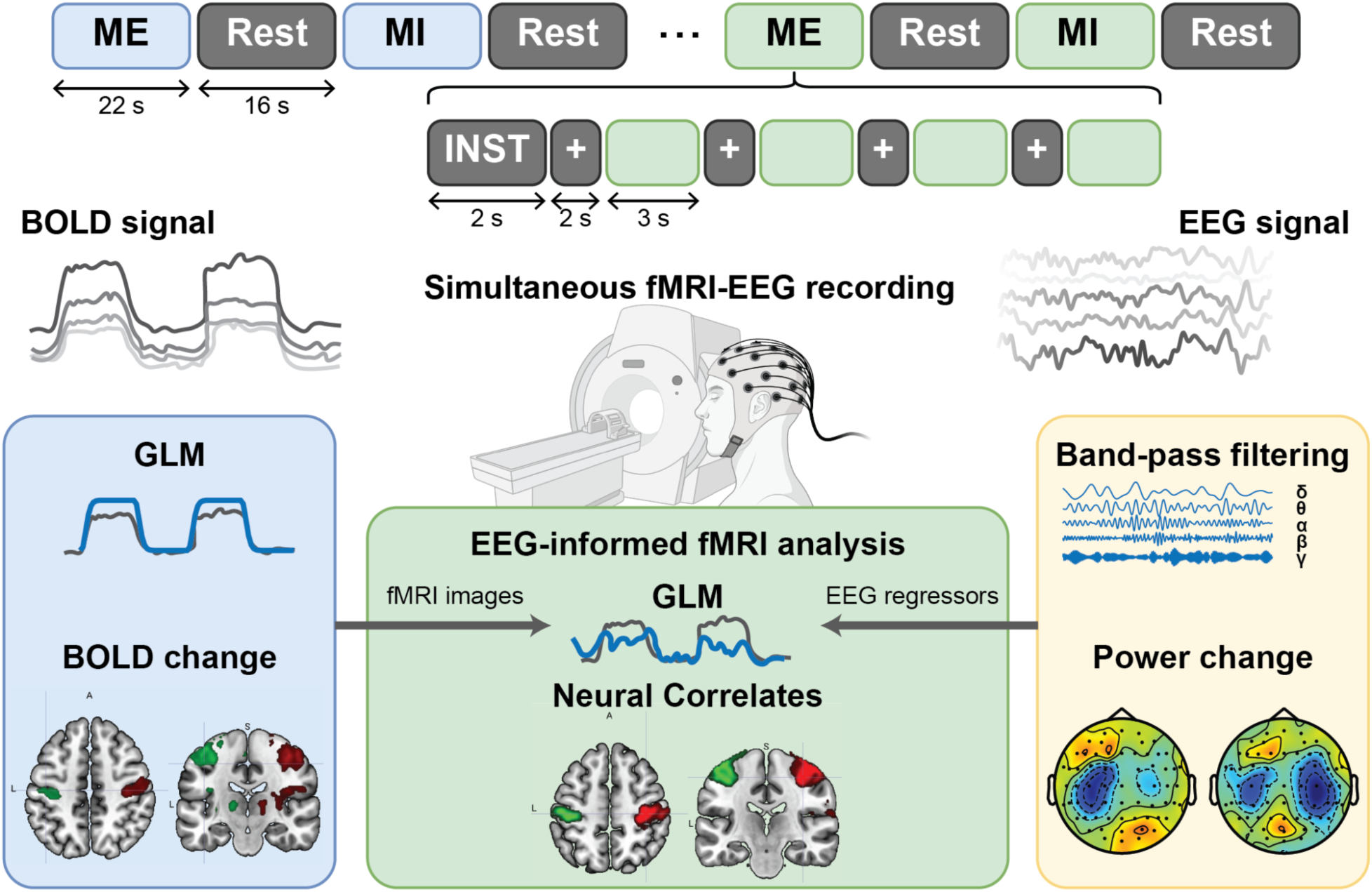
Overview of the experimental design and data processing strategy. EEG and fMRI data were simultaneously recorded during the session. The analysis included both separate and integrated approaches for these modalities.

### 2.3 EEG-fMRI Data Acquisition

EEG was simultaneously recorded during fMRI acquisitions using an MR-compatible EEG system (BrainAmp MR plus, Brain Products GmbH, Gilching, Germany). The EEG cap (BrainCap MR, EasyCap GmbH, Breitbrunn, Germany) included 63 scalp electrodes distributed according to the 10–20 system. One additional electrocardiographic (ECG) electrode was placed on the participants’ chest. All EEG signals were relative to a reference located in FCz, with the ground in correspondence of AFz. The sampling frequency of EEG and ECG data was set at 5,000 Hz. The impedance at each electrode was kept below 10 kΩ.

The MRI acquisition was performed on a 3T MRI scanner (Siemens Magnetom Prisma, Erlagen, Germany) with a 64-channel head coil at the CMU-Pitt BRIDGE Center (RRID:SCR_023356). Each subject underwent one MRI session composed of a structural MRI and three consecutive task fMRI scans. Brain anatomy was acquired with a T1-weighted image with 3D magnetization-prepared rapid gradient echo (MPRAGE) sequence (matrix size: 256×256×192 mm; resolution:1×1×1 mm^3^; repetition time (TR) = 2300 ms, echo time (TE) = 1.9 ms; inversion time = 900 ms; flip angle (FA)= 9°). For each fMRI acquisition, 460 volumes were acquired with a 2D echo planar imaging sequence with spin echo (matrix size: 212×212×144 mm; resolution:2×2×2 mm^3^; TR = 2s; TE = 30.0ms; FA = 79°; 72 axial slices), for a total time of 15 min 33 sec.

### 2.4 EEG Analysis

EEG data recorded in the MRI scanner were initially processed in BrainVision Analyzer (Brain Products GmbH, Gilching, Germany) for gradient artifact (GA) and ballistocardiogram (BCG) artifact correction. The GA was removed from each channel by subtracting a template generated from a sliding average of 21 blocks of the artifact (Allen et al., 2000). The cleaned EEG and ECG data were subjected to a bandpass filtering of 0.1 and 70 Hz and 0.5 and 30 Hz, respectively, with a notch filter of 60 Hz, and downsampled to 250 Hz. The BCG artifact was then addressed by semi-automatically identifying the R peaks on the ECG and subtracting a 21-block sliding window average artifact template from the data (Allen et al., 1998; Allen et al., 2000). Bad intervals were identified through visual inspection and eye movements, and any remaining BCG artifacts were removed by applying an extended Infomax Independent Component Analysis (ICA) (Maggioni et al., 2014).

Following offline correction for MRI-induced artifacts, data were further cleaned using the FieldTrip toolbox (Oostenveld et al., 2011) and custom MATLAB scripts (MathWorks Inc., MA, USA). The EEG data were bandpass filtered between 0.2 and 40 Hz. The continuous EEG signal was then segmented into trials, each spanning from 2 seconds before trial onset to 1 second after the end of the trial. ICA was applied to the segmented data to remove residual artifacts. Trials with a standard deviation exceeding 100 µV were excluded from further analysis.

Single-trial time-frequency representations (TFRs) of the EEG data were calculated using Morlet wavelets. Baseline power was determined as the average power at each frequency within the 1-second to 0.1-second window preceding trial onset. Event-related desynchronization (ERD) was calculated for each time-frequency pair as the percentage change relative to baseline power, using the formula: ERD(*t,f,c*) = (P(*t, f,c*) − R(*f,c*)) / R(*f,c*) × 100%, where R(*f,c*) denotes the average baseline power at frequency *f* and channel *c*, and P(*t,f,c*) represents the power at time point *t*, frequency *f*, and channel *c*. The single-trial alpha band ERD was quantified by averaging the ERD from 0.5 seconds after trial onset until the end of the trial within the alpha frequency band (8 - 13.5 Hz).

### 2.5 fMRI Analysis

fMRI analyses were performed using the Statistical Parametric Mapping (SPM) MATLAB toolbox (http://www.fil.ion.ucl.ac.uk/spm/, version 12) and custom MATLAB scripts.

#### 2.5.1 fMRI preprocessing

To obtain clean fMRI data, fMRI volumes were spatially realigned to the first volume to reduce head motion artifacts, distortion artifacts were corrected using the topup tool of the FMRIB Software Library (FSL) software (version 6) (Jenkinson et al., 2012), fMRI volumes were then co-registered with the T1-weighted image, normalized according to Montreal Neurological Institute (MNI) standard space, and spatially smoothed using a 4 mm size full width half maximum 3D Gaussian kernel. The profile of head motion was evaluated by calculating the framewise displacement (FD) ( Power et al., 2012). One subject was discarded for excessive head motion, showing a FD higher than three times the group standard deviation (FD mean = 0.32, SD

= 0.21).

#### 2.5.2. Unimodal block fMRI analysis

A block fMRI analysis was performed using MATLAB custom scripts and SPM to localize the hemodynamics correlates of the task performed inside the scanner. For each participant, a first-level General Linear Model (GLM) was applied to the fMRI images. The BOLD activity at each voxel was model ed as a linear combination of task and movement regressors - the parameters of head rotation (N=3) and translation (N=3) obtained from the fMRI preprocessing steps - which served as regressors of interest and confounders, respectively. Each task regressor, representing the model ed BOLD response for a specific task type, was obtained by the convolution of the onset of the trials and the canonical hemodynamic response function (HRF), downsampled to match the fMRI sampling frequency (fs = 0.5 Hz), and high-pass temporally filtered with a 128 s cut-off. A scan-based model was used to account for the three consecutive scans performed during the acquisition. Therefore, each scan was model ed separately in the GLM assuming consistency of task-related effects among different sessions. Hemodynamic activation of different tasks was firstly assessed at subject level using t-statistic, obtaining first-level fMRI contrasts maps for the following t-contrasts: “ME RH > ME LH”, “ME LH > ME RH”, “MI RH > MI LH”, “MI LH > MI RH”. Subsequently, first-level fMRI contrast maps were entered into a second-level random-effects design. The GLM analysis incorporated the contrast maps as dependent variables while using age, sex, and handedness as covariates. Following GLM estimation, we applied t-contrasts to evaluate group-level responses (p < 0.001, k ≥ 15 voxels). We used the Automated Anatomical Atlas 3 (AAL3) (Rolls et al., 2020) to identify the anatomical locations of significant clusters.

### 2.6 Cross-Modal Correlation

To examine the relationship between EEG and fMRI signals, a correlation analysis was conducted between EEG power changes and BOLD signal changes during MI tasks. The regions of interest (ROI) for fMRI responses were defined as the contralateral sensorimotor region, based on literature, specifically the precentral and postcentral gyri, based on the AAL3 (Rolls et al., 2020). The mean t-statistic from the first-level fMRI analysis within the ROI after thresholding at p < 0.001 was extracted using MarsBaR (Brett et al., 2002). For each subject, the average percentage changes in high-alpha band EEG power (10.5 - 13.5 Hz) was extracted from channels C3 and C4 for right-hand and left-hand MI tasks, respectively. A linear regression was then conducted to assess the relationship between contralateral ERD and the contralateral t-statistics from the BOLD response at the group level (N=17).

### 2.7 EEG-Informed fMRI Analysis

The data-driven investigation of hemodynamic correlates of the neural activity was conducted using an EEG-informed fMRI framework by exploiting the spectral content over time of EEG signal.

Following the removal of MR-induced artifacts, the processing of continuous data for EEG-informed fMRI analysis was performed using FieldTrip toolbox (Oostenveld et al., 2011) and custom MATLAB scripts (MathWorks Inc., MA, USA). Bad trials, identified using the same methodology described above for the EEG epoched signal, were replaced by the continuous signal with the average of the correspondent trial type signals. The time-frequency content of the continuous signal was extracted by computing the Continuous Wavelet Transform (CWT) using the Morlet wavelet family in the frequency range 1 - 40 Hz with a step of 0.5 Hz and a central frequency of 0.8125 Hz. The time-varying power of the signal was calculated as the squared module of the CWT and averaged within the frequency band of interest. For our study, given that the experimental protocol involved right-hand and left-hand motor tasks, we focused on alpha frequency band (8 - 13.5 Hz) and selected the central electrodes above the hand knobs within the motor cortices (C3 and C4), following the approach of previous studies (Laufs et al., 2003; Maggioni et al., 2016; Sclocco et al., 2014), that showed a clear ERD in our participants during the MI tasks. In this framework, as shown in Fig. 2, time-frequency content from both electrodes is simultaneously fed into the model using a data-driven approach, showing the model’s capability to discriminate between right-hand and left-hand movements and imaginations without relying on priori information about the sensor placement or the task onset time.

**Figure 2.**
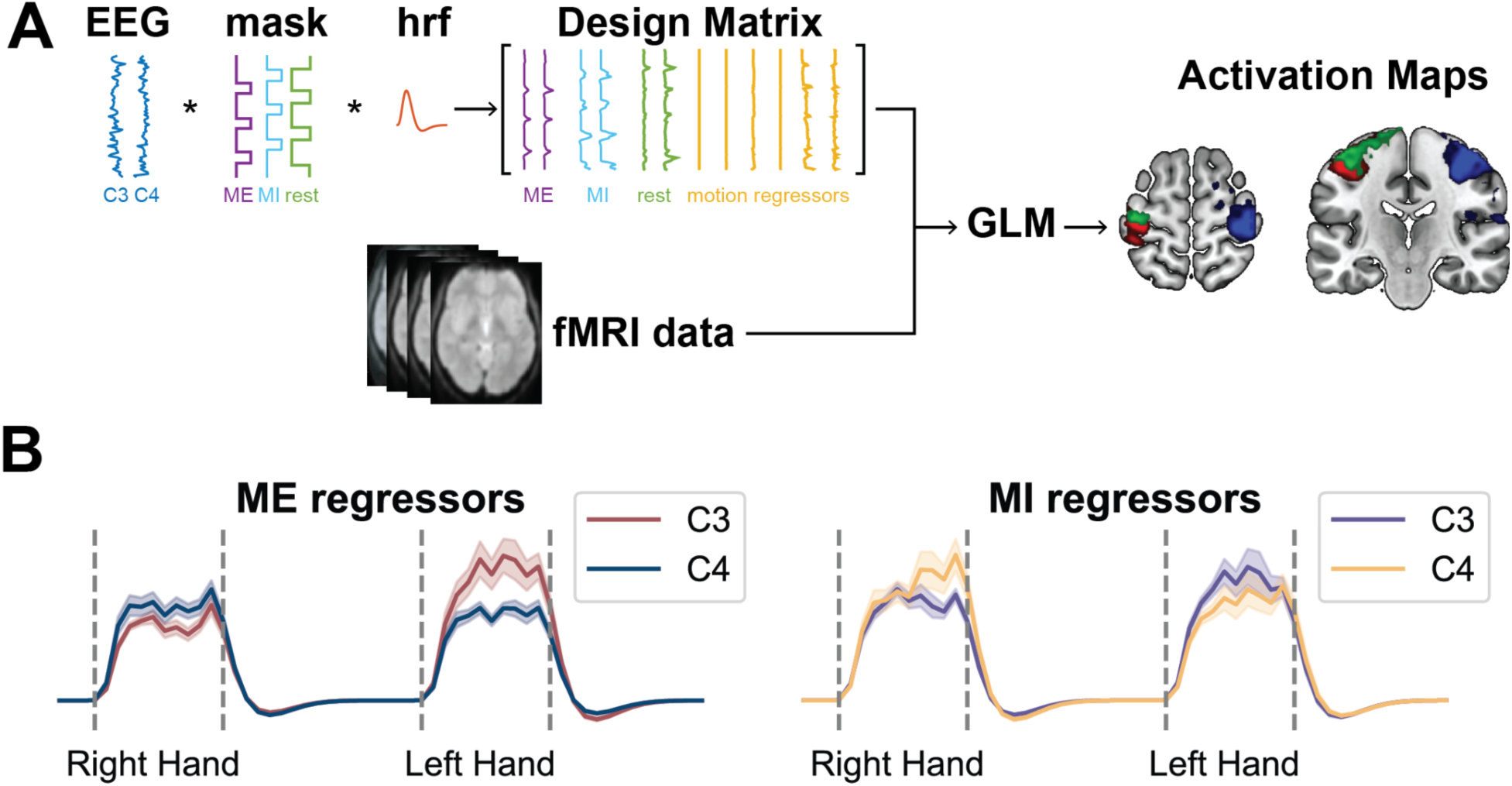
**A:** Overview of the EEG-informed fMRI analysis pipeline. The time-frequency signals in alpha band (8 - 13.5 Hz) were separated into three regressors (ME, MI, and rest) based on the experiment structure. These regressors were convolved with the hemodynamic response function, downsampled to match the fMRI sampling frequency, and high-pass filtered. Subject-specific design matrices were built with three regressors each for C3 and C4, along with movement parameters. A fixed-effects General Linear Model (GLM) was applied at the group level, with t-statistics used to extract contrasts of interest. **B:** Example regressors for ME and MI tasks from individual subjects.

The regressors of interest of the EEG-informed fMRI analysis were built from the continuous time-frequency signal of C3 and C4 in the alpha frequency band previously calculated. The following analyses were performed on a subsample of subjects that showed a clear activation pattern in the contralateral sensorimotor region during the MI tasks. The group of subjects was identified calculating the precision of the single-subject unimodal fMRI results with respect to the contralateral sensorimotor region, using the following formula: TP/(TP+FP) x 100%, where TP (true positive) represents the number of voxels of the unimodal fMRI activation map within the corresponding sensorimotor region, whereas FP (false positive) denotes the number of voxels outside the target region. The average precision of “MI RH > MI LH”, “MI LH > MI RH” contrasts were calculated and only subjects with a precision higher than 25% were retained for the following analyses. Eight out of seventeen participants (two males, six females; aged 23.13 ± 1.46 years; six right-handed) were included in the following analyses.

To investigate the hemodynamic correlates of ME and MI activity, the signals were first separated into three different regressors reflecting the experiment structure (ME, MI, and rest) by masking the time-frequency signal with the corresponding squared waves indicating the duration of the three experiment conditions. Afterwards, the signal was convolved with the HRF, downsampled to match the fMRI sampling frequency (fs = 0.5 Hz), and high-pass temporally filtered with a 128 s cut-off. For each subject, the regressors of interest, three for C3 and three for C4, were entered along with the movement parameters in a scan -based model design matrix. The group-level results were evaluated through a fixed-effects GLM model where the design matrix included the design matrices of all subjects. Beta coefficients estimated for C3 and C4 regressors for task conditions of interest (ME and MI) were compared by applying t-statistics and obtaining contrasts maps for the following t-contrasts: “C3 < C4 - ME”, “C4 < C3 - ME”, “C3 < C4 - MI”, “C4 < C3 - MI”. T-contrasts maps were used to assess significant results (p < 0.001, k ≥ 15 voxels) and the anatomical locations of significant clusters were identified with the AAL3 (Rolls et al., 2020). The hypothesis underlying this analysis is that the BOLD signal of the regions involved in a movement execution or imagery task are more negatively correlated with the alpha power of the sensor directly above that region with respect to the alpha power of the sensor on the contralateral side. Therefore, for example, when investigating the t-contrast “C3 < C4 – ME” we expect to find regions involved in the right-hand motor execution task, such as the left sensorimotor cortex underlying C3, with the model not having information a priori on electrode positions or the timing of task execution.

The degree of accuracy of the group-level EEG-informed fMRI analysis was evaluated by comparing the results with the unimodal fMRI analysis. In order to quantitatively compare the results, group-level fMRI results were calculated with a fixed-effects GLM model over the best performing subjects, as applied for the EEG-informed fMRI analysis. The quantitative overlap was assessed with the dice coefficient (DC), precision, and recall indices calculated over the binarized images (p < 0.001, k ≥ 15 voxels). DC gives an indication of the amount of overlap between the two images, being sensitive to the dimension of both images, and was calculated with the formula: 2 x TP / ( 2 x TP + FP + FN) x 100%; precision provides information on the overlap between the two images taking into account the extent of the EEG-informed fMRI image with the following formula: TP/(TP+FP) x 100%; recall represents how much the overlap between the two images is within the fMRI image and it was calculated using the formula: TP/(TP+FN) x 100%. In the formula, TP represents the number of voxels of EEG-informed fMRI that fall within the fMRI region, FP is the number of EEG-informed fMRI voxels that fall outside the fMRI activated regions, and FN (false negative) represents the number of voxels of fMRI that are not overlapped with the EEG-informed fMRI.

## 3. Results

The experimental paradigm and analytical framework are depicted in Fig. 1. EEG and fMRI data were recorded simultaneously during the session, in which participants performed alternating ME and MI tasks involving hand and foot movements. A mixed block- and trial-based design, adapted from Yuan et al. (2010), was implemented to facilitate integrated EEG and fMRI analysis. Task-induced electrophysiological activity was quantified by extracting ERD values from EEG signals, while changes in BOLD signals captured task-related hemodynamic responses. The covariation between task-specific EEG and BOLD signals was examined through a linear regression model. An EEG-informed fMRI analysis was conducted to further elucidate neurovascular coupling, modeling the interaction between EEG rhythms and BOLD responses.

### 3.1 In-scanner EEG reveals contralateral ERD during hand motor imagery

Decreases in alpha and beta band power over contralateral sensorimotor areas were observed during hand MI tasks in the offline-corrected EEG data collected simultaneously with fMRI scanning (Fig. 3A). Spatial distributions illustrating the relative changes in alpha band power (8 - 13.5 Hz) are displayed in Fig. 3B. The most pronounced suppression was localized around channel C3 during right-hand MI tasks and around channel C4 during left-hand MI tasks. Although power decreases were also present on the ipsilateral side of the brain, they were smaller in magnitude and extent compared to those on the contralateral side. The average contralateral ERD was -16.18% (SD = 31.62%) for right-hand MI and -18.58% (SD = 18.36%) for left-hand MI.

**Figure 3.**
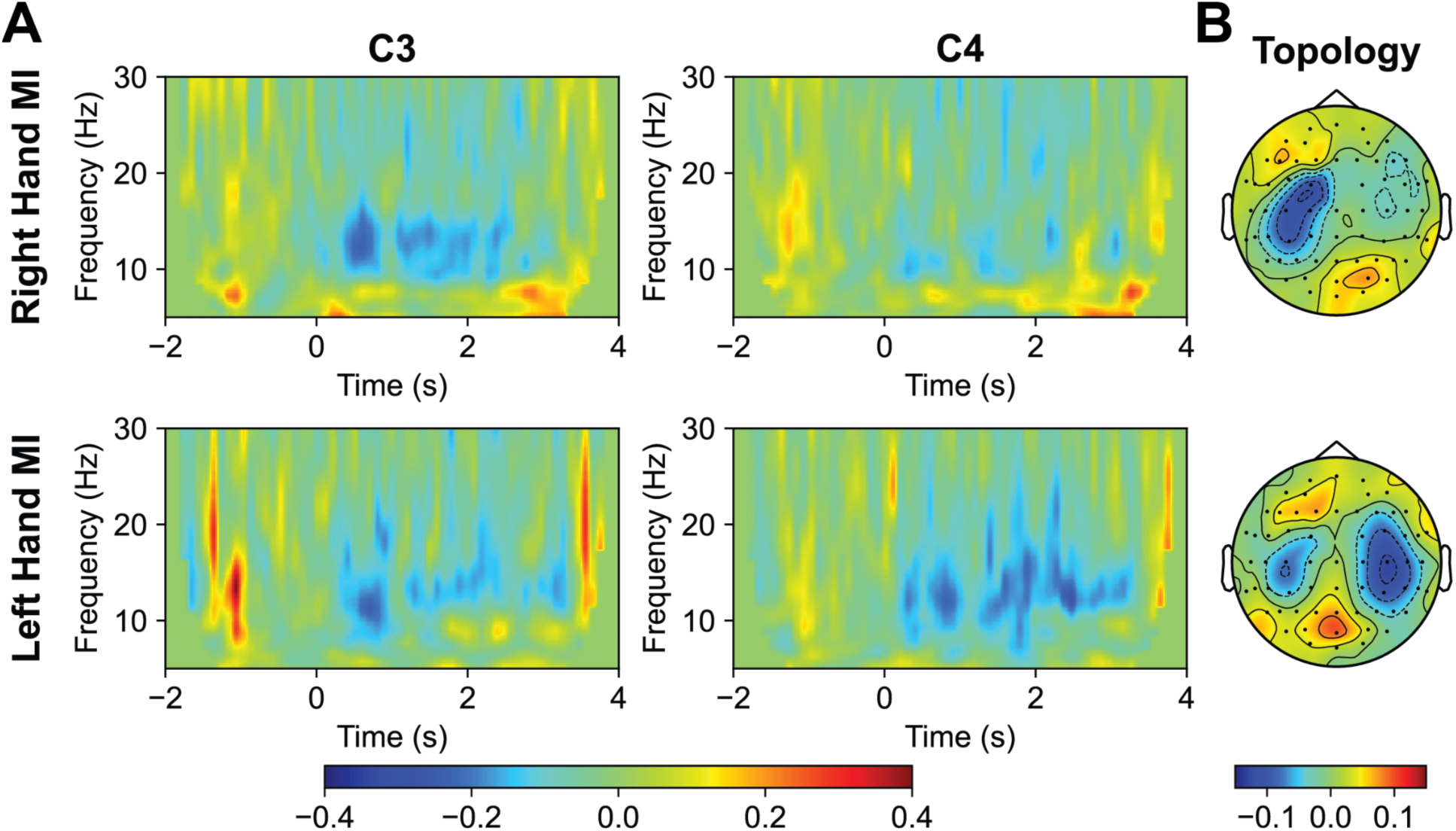
**A:** Group-averaged time-frequency maps for C3 and C4 during right-hand MI/left-hand MI (n=17). **B:** Group-averaged alpha ERD topology for right-hand MI/left-hand MI (8 – 13.5 Hz) (n=17).

### 3.2 Unimodal fMRI analysis shows task-induced activations

Fig. 4A - 4D shows the group-level BOLD activation surface maps related to the performed ME and MI tasks, whereas Tables S1 and S2 report the detailed analysis results. When the BOLD activation for ME and MI of one hand was compared to that of the opposite hand, increases in BOLD signal were observed in the contralateral sensorimotor areas during both executed and imagined hand movements. Notably, motor execution elicited a substantially stronger hemodynamic response than motor imagery. The ME-related activations included regions such as the contralateral primary motor cortex (BA4), primary somatosensory cortex (BA3), supplementary motor area (SMA, BA6), thalamus, putamen, and insular cortex (BA13), and ipsilateral activations in the cerebellum. Similarly, the MI-induced BOLD activities included areas like contralateral primary motor cortex (BA4) and primary somatosensory cortex (BA3). Moreover, when comparing right-hand MI with left-hand MI, BOLD activations were found in the nucleus accumbens, supramarginal (BA40), middle cingulate regions, secondary visual cortex (BA18), and right cerebellum, whereas left-hand MI showed higher activation in the right putamen when compared with right-hand MI.

**Figure 4.**
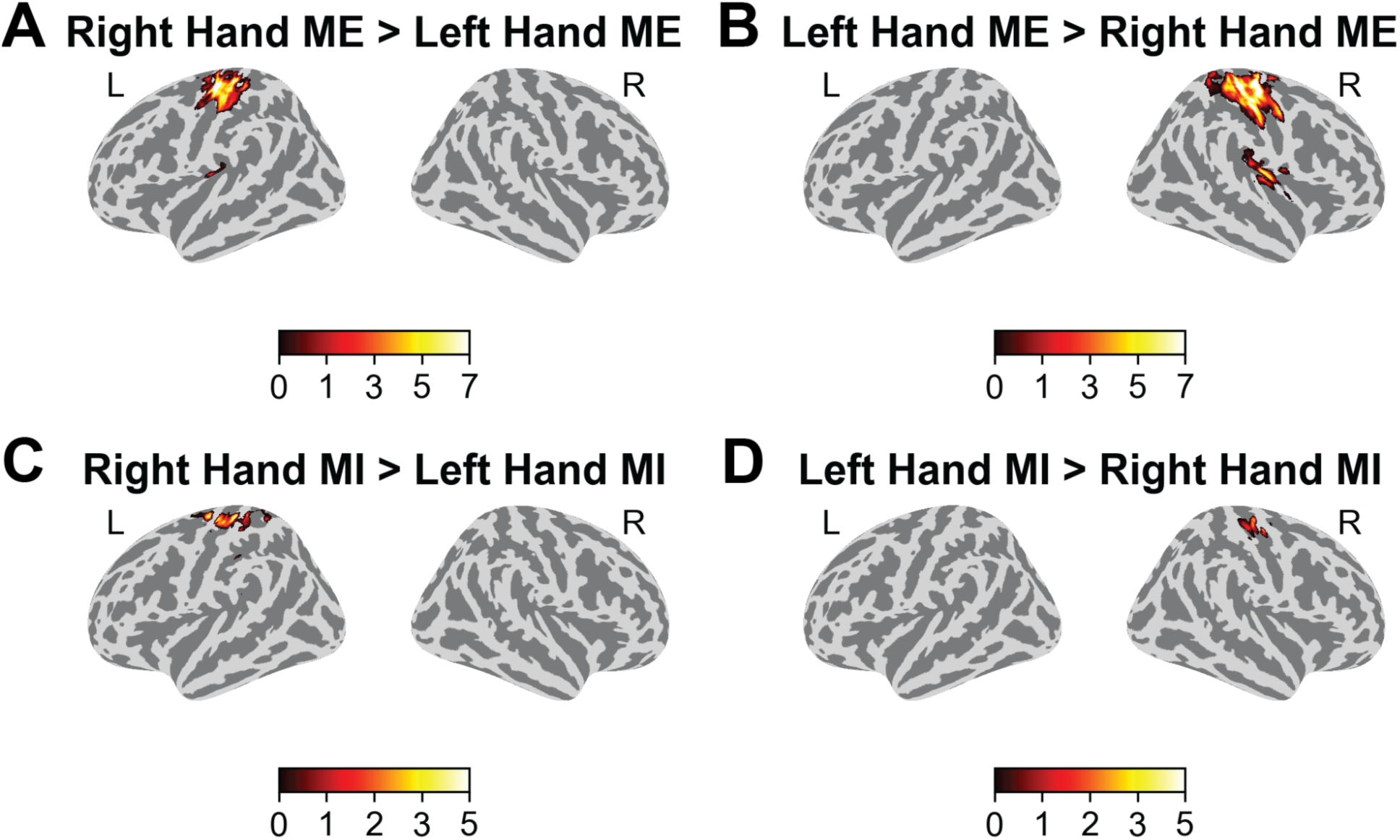
**A:** Group-averaged t values of the contrast right-hand ME > left-hand ME (p < 0.001, k ≥ 15 voxels) (n = 17). **B:** Group-averaged t values of the contrast left-hand ME > right-hand ME (p < 0.001, k ≥ 15 voxels) (n = 17). **C:** Group-averaged t values of the contrast right-hand MI > left-hand MI (p < 0.001, k ≥ 15 voxels) (n = 17). **D:** Group-averaged t values of the contrast left-hand MI > right-hand MI (p < 0.001, k ≥ 15 voxels) (n = 17).

### 3.3 ERD power change negatively correlates with BOLD activity

Fig. 5A presents the group-averaged relative changes in EEG alpha band power at c hannels C3 and C4, while Fig. 5B illustrates the group-averaged t-statistics representing task-induced BOLD changes within the sensorimotor region. We found decreases in EEG alpha band power were correlated with increases in BOLD responses on the same side of the brain. During both left and right-hand MI tasks, the contralateral hemisphere exhibited EEG power suppression coupled with heightened BOLD activation. To further investigate the relationship between these two modalities, we conducted a linear regression analysis between contralateral ERD and contralateral t-values from the BOLD response, which indicated a negative covariation between EEG and BOLD changes (r = -0.49, p < 0.05, Fig. 5C).

**Figure 5.**
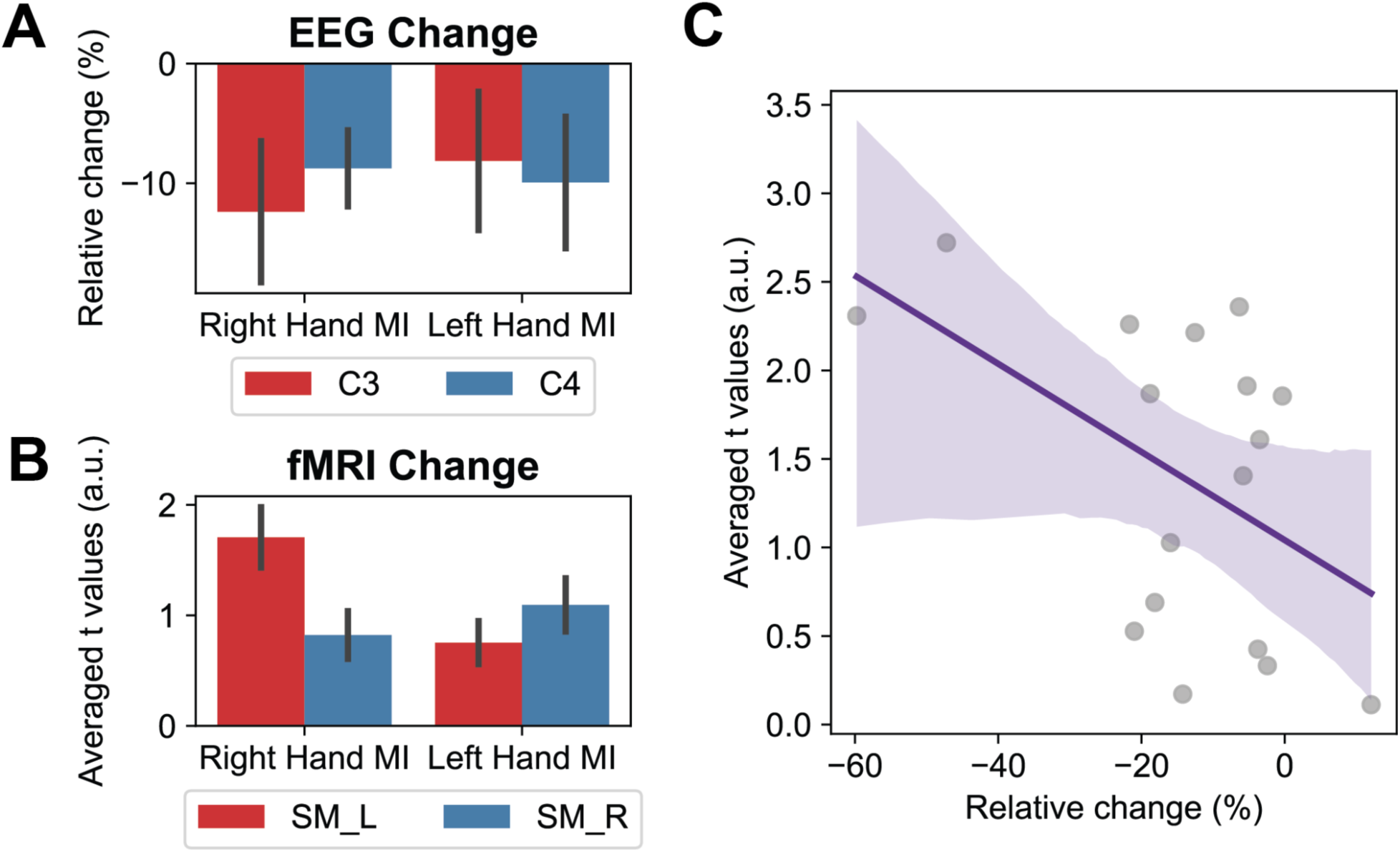
**A:** Group-averaged high-alpha ERD (10.5-13.5 Hz) (n=17). **B:** Group-averaged t values within the ROI after thresholded by p < 0.001 (SM_L: left sensorimotor region; SM_R: right sensorimotor region, including precentral and postcentral gyrus) (n=17). **C:** Correlation between contralateral ERD and contralateral averaged t values within the contralateral sensorimotor region after thresholded by p < 0.001 (r = -0.49, p-value < 0.05) (n=17).

### 3.4 EEG-informed fMRI analysis shows whole-brain hemodynamic correlates of EEG alpha power

Fig. 6 displays the group-level BOLD activation maps of the EEG-informed fMRI analysis, whereas Table 1 reports the detailed analysis results. Consistent with our hypothesis, increased activation in the sensorimotor region was observed when the alpha power of the ipsilateral electrode of interest was lower than that of the contralateral electrode of interest, for both MI and ME. In ME-related “C3 < C4” contrast (Fig. 6B), the analysis highlights the activation of the left primary motor cortex (BA4), left primary somatosensory cortex (BA3), right angular gyrus (BA39), right temporal area (BA21, 22), right cerebellum, and right middle frontal gyrus (BA8,9). Oppositely, in the ME-related “C4 < C3” contrast (Fig. 6C), hemodynamic correlates reflect regions active during the execution of the left-hand movement, as the right primary motor cortex (BA4), right primary somatosensory cortex (BA3), left inferior frontal gyrus (BA47), right insular cortex (BA13), left cerebellum, and right SMA (BA6). Finally, in MI-related contrast “C3 < C4” (Fig. 6A), the analysis revealed an increase of BOLD signal in the left primary motor cortex (BA4), left primary somatosensory cortex (BA3), visual cortex (BA17, 18, 19), and left superior frontal area (BA9). No significant results were found for the MI-related contrast “C4 < C3”.

**Figure 6.**
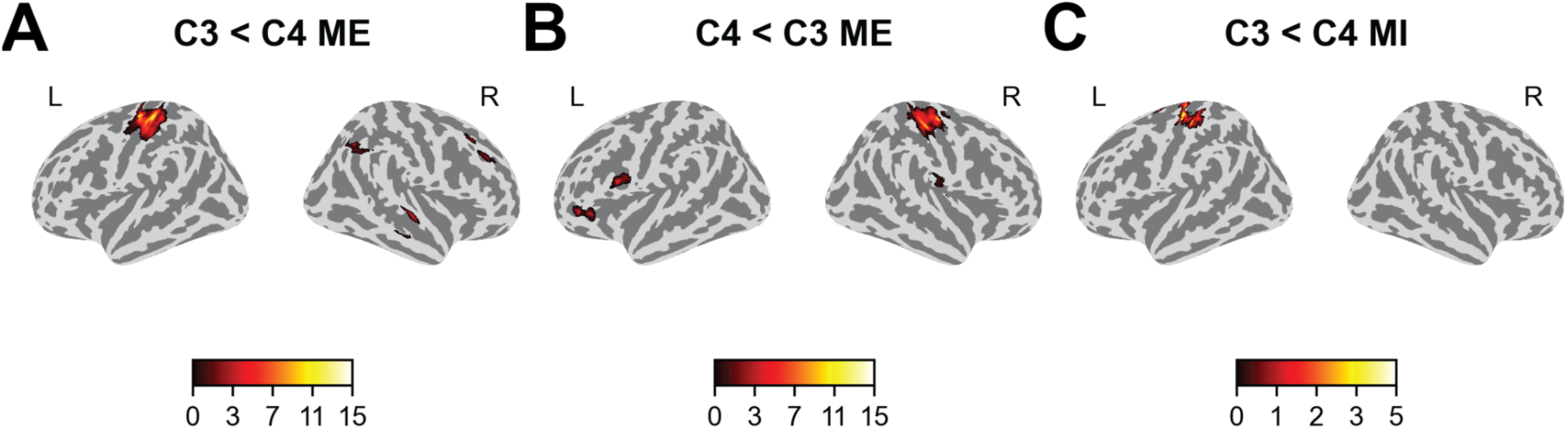
**A:** Group-averaged t values showing neural correlates of alpha rhythms in channel C3 during ME tasks (p < 0.001, k ≥ 15 voxels) (n = 8). **B:** group-averaged t values showing neural correlates of alpha rhythms in channel C4 during ME tasks (p < 0.001, k ≥ 15 voxels) (n = 8). **C:** Group-averaged t values showing neural correlates of alpha rhythms in channel C3 during MI tasks ( p < 0.001, k ≥ 15 voxels) (n = 8).

**Table 1.** Group-level EEG-informed fMRI activations using alpha-power regressors for motor execution (ME) and motor imagination (MI) using a fixed-effects model (p < 0.001, k ≥ 15 voxels) (n = 8). Labeling performed with AAL3 atlas.

**C3 < C4 - ME**
| T-value<br>($p < 0.001$, $k > 15$) | Cluster-size<br>[# voxels] | Peak coordinates<br>(xyz) [mm] | Atlas regions (%) |
| --- | --- | --- | --- |
| 16.0 | 1007 | -38 -26 62 | Left Postcentral gyrus (80), Left Precentral gyrus (18), Left Superior parietal gyrus (2) |
| 5.6 | 65 | 46 -60 50 | Right Angular gyrus (45), Right Superior parietal gyrus (34), Right Inferior parietal gyrus-excluding supramarginal and angular gyri (22) |
| 5.6 | 65 | 64 -16 -4 | Right Superior temporal gyrus (51), Right Middle temporal gyrus (49) |
| 4.9 | 81 | 28 -40 -24 | Right Lobule IV-V of cerebellar hemisphere (85), Right Fusiform gyrus (15) |
| 4.7 | 32 | 32 28 32 | Right Middle frontal gyrus (81), Right Inferior frontal gyrus-triangular part (6) |
| 4.3 | 44 | 32 30 54 | Right Middle frontal gyrus (86), Right Superior frontal gyrus-dorsolateral (14) |

**C4 < C3 - ME**
| T-value<br>($p < 0.001$, $k > 15$) | Cluster-size<br>[# voxels] | Peak coordinates<br>(xyz) [mm] | Atlas regions (%) |
| --- | --- | --- | --- |
| 14.0 | 836 | 44 -24 64 | Right Postcentral gyrus (71), Right Precentral gyrus (24) |
| 6.9 | 86 | -40 36 -4 | Left Inferior frontal gyrus-triangular part (59), Left IFG pars orbitalis (41) |
| 5.4 | 55 | -46 8 16 | Left Precentral gyrus (65), Left Inferior frontal gyrus-opercular part (35) |
| 5.1 | 36 | -86 7 6 | Left Supplementary motor area (53), Left Superior frontal gyrus-dorsolateral (36) |
| 4.8 | 34 | 52 -16 20 | Right Rolandic operculum (94), Right Insula (3), Right Superior temporal gyrus (3) |
| 4.4 | 72 | -16 -52 -18 | Left Lobule IV-V of cerebellar hemisphere (92), Left Lobule VI of cerebellar hemisphere (7), Left Fusiform gyrus (1) |
| 4.3 | 24 | 12 -10 54 | Right Supplementary motor area (100) |

**C3 < C4 - MI**
| T-value<br>($p < 0.001$, $k > 15$) | Cluster-size<br>[# voxels] | Peak coordinates<br>(xyz) [mm] | Atlas regions (%) |
| --- | --- | --- | --- |
| 5.5 | 541 | -12 -68 -4 | Left Lingual gyrus (49), Left Calcarine fissure and surrounding cortex (43), Left Cuneus (4), Right Calcarine fissure and surrounding cortex (3) |
| 4.6 | 337 | -14 -22 78 | Left Precentral gyrus (48), Left Postcentral gyrus (42), Left Paracentral lobule (7) |
| 3.6 | 19 | -16 -2 62 | Left Superior frontal gyrus-dorsolateral (89) |

The quantitative comparison between EEG-informed fMRI results and unimodal fMRI results was performed on results obtained from the same cohort. fMRI results on the sub-samples were in line with the results obtained at the group level (Table S3, Table S4). The comparison showed high overlap between “C3 < C4” ME and MI contrasts with right-hand execution and imagination fMRI maps, respectively, and that “C4 < C3” ME contrast was overlaid with left-hand execution fMRI map. Specifically, the alpha ME regressors (Fig. 7A, B) showed hemodynamic correlates in the sensorimotor regions (BA3,4,6), cerebellum, and insular cortex (BA13), showing DC equal to 0.33, precision of 0.84, and recall of 0.20 for right-hand related results, and DC equal of 0.24, precision equal to 0.84, and recall of 0.14 for left-hand related maps. Lastly, MI-related right-hand imagination (Fig. 7C, 7D) activations overlapped over the left sensorimotor cortex (BA3,4) and left superior frontal gyrus (BA9) with a DC of 0.10, precision equal to 0.36, and recall of 0.06. When restricting the EEG-informed fMRI and fMRI results to contralateral sensorimotor regions, greater overlap was observed in activated areas, while the EEG-informed fMRI model provided more focalized results (Fig. S1). For right-hand ME, the precision was 1.00 (DC = 0.42, recall = 0.27), for left-hand ME, precision was 1.00 (DC = 0.52, recall = 0.35), and for right-hand MI, precision was 0.98 (DC = 0.19, recall = 0.10).

**Figure 7.**
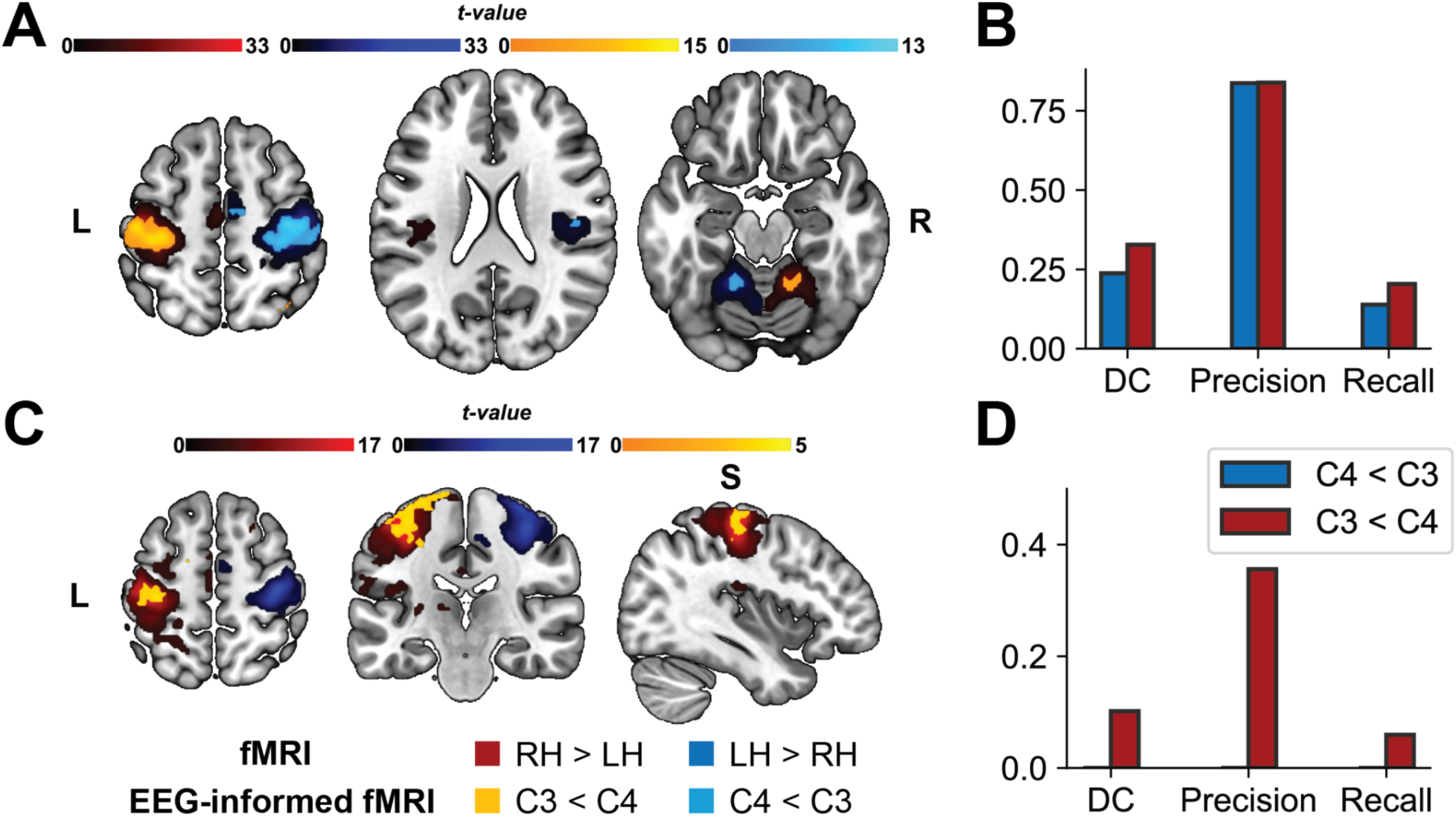
**A:** Group-level fMRI statistical maps for ME conditions, showing the overlap between fMRI analysis results (RH > LH; LH > RH) and EEG-informed fMRI analysis results (C3 < C4; C4 < C3) (n = 8). Axial slices with MNI coordinates at z = 56, z = 22, and z = -18 are displayed from left to right. **B:** Quantitative overlap between fMRI analysis and EEG-informed fMRI analysis results for ME tasks, presented as dice coefficient (DC), precision, and recall. **C:** Group-level fMRI statistical maps for MI conditions, showing the overlap between fMRI analysis results (RH > LH; LH > RH) and EEG-informed fMRI analysis results (C3 < C4; C4 < C3) (n = 8) [x = -38, y = -22, z = 58]. **D:** Quantitative overlap between fMRI analysis and EEG-informed fMRI analysis results for MI tasks, presented as dice coefficient (DC), precision, and recall.

## 4. Discussion

In this study, we have examined the neural patterns associated with various motor execution and motor imagery tasks using simultaneous EEG and fMRI recordings. Correlation analysis of the two modalities confirmed the inverse relationship between task-related EEG power changes in the alpha frequency band and hemodynamic responses during motor tasks. For the first time, we applied EEG-informed fMRI analysis to tasks involving both ME and MI, successfully distinguishing ME and MI activities across multiple task conditions through a unified model. Our findings reveal continuous, temporal covariation between task-induced EEG power changes in the sensorimotor region and BOLD activations in both colocalized and coactivated regions. The substantial overlap between the EEG-informed fMRI results and block fMRI results further supports the robustness and reliability of EEG-informed fMRI in capturing electrophysiological-hemodynamic coupling across diverse task conditions, even in the absence of prior information about these conditions.

The neural patterns underlying ME and MI activities have been studied using noninvasive methods, including EEG and fMRI. In our results, we observed bilateral suppression of alpha power in EEG during MI, with a stronger and more widespread ERD on the contralateral side. This finding aligns with prior research on ERD during MI activities, which consistently reports contralateral ERD patterns in the alpha and beta bands, while EEG changes on the ipsilateral side tend to vary across subjects and studies (Pfurtscheller & Lopes da Silva, 1999; Wang et al., 2010; Wang et al., 2024). Voluntary and imaginary movements involve the activation of similar networks, coordinating sensorimotor integration and motor execution, such as motor, pre-motor, parietal cortex, and subcortical structures, such as the cerebellum, thalamus, and basal ganglia (Hardwick et al., 2018; Henschke et al., 2023). During actual movements, the SMA, a region involved in motor planning (Fried et al., 2011), receives input from the lateral prefrontal cortex and deep structures, such as basal ganglia, thalamus, and cerebellum (Rozzi et al., 2017; Sakai et al., 2013). Besides these regions, our results also show the involvement of the posterior part of the insula in motor execution tasks, a region specifically involved in interoception, perceptual self-awareness, multimodal signal processing, and autonomic control (Benarroch, 2019), which may play a role in processing the effects of gravity by integrating sensory feedback information, especially in its posterior region (Rousseau et al., 2021). The regions activated for ME and MI showed significant overlap, confirming the similarity in neurophysiological mechanisms underlying these two activities (Decety, 1996). Compared to ME, the BOLD responses induced by MI were of lower intensity and more limited in extent, which is consistent with previous findings (Decety, 1996; Hanakawa et al., 2003; Yuan et al., 2010). Besides common activations, right-hand MI showed higher activation in the middle cingulate gyrus (MCG) and secondary visual cortex. MCG is known to be involved in coordinating motor planning and execution, including imagined movement (Hardwick et al., 2018; Wang et al., 2023). Since motor imagery can involve visual streams (Glover & Baran, 2017; Jiang et al., 2015), the higher activation of the secondary visual cortex could be related to participants relying more on vivid visual imagery for the right-hand rather than the left one.

Previous studies have investigated the relationship between BOLD signals and EEG rhythms across various contexts. Positive correlations have been observed between BOLD signals and delta band power in resting states (Portnova et al., 2018). Beta power, by contrast, negatively correlates with BOLD activity in the left inferior prefrontal cortex during memory formation (Hanslmayr et al., 2011). Gamma band oscillations in EEG have shown positive associations with BOLD activation in the visual, auditory, and motor cortices (Brookes et al., 2005; Mulert et al., 2010; Uji et al., 2018). These findings suggest a complex relationship between the hemodynamic response and EEG fluctuations at different frequency bands. Studies have reported the colocalization and the negative covariation between EEG alpha power and fMRI BOLD changes during motor tasks (Sclocco et al., 2014; Yuan et al., 2010; Zich et al., 2015). Interestingly, similar inverse correlations between alpha power and BOLD signals have also been observed in other cortical regions under diverse cognitive conditions. For example, negative correlations between alpha rhythms and BOLD responses in the visual cortex are well-documented (Bareither et al., 2014; Becker et al., 2015). Additionally, Tagliazucchi et al. (2012) found that increased alpha power corresponds with decreased fMRI BOLD connectivity. Our findings further support this inverse relationship between EEG alpha rhythms and hemodynamic responses during motor imagery through a correlation analysis, consistent with the understanding that alpha rhythms often signify reduced cortical activity as an idling state. In the correlation analysis, the high alpha frequency band (10.5– 13.5 Hz) was analyzed instead of the broader alpha band (8–13.5 Hz) because the high alpha component is believed to offer a more nuanced insight into motor cortex activation, which therefore better distinguished individuals with various MI expertise (Pfurtscheller et al., 2000; Zabielska-Mendyk et al., 2018).

We have confirmed and extended the previous findings on the correlation between EEG power change and BOLD changes during motor activities through joint EEG-fMRI analysis. In recent years, EEG-fMRI has gained considerable interest as an approach that allows integrating the high temporal resolution of EEG with the spatial precision provided by fMRI. In previous studies investigating correlations between electrophysiological and hemodynamics features during motor imagery exploiting simultaneous EEG and fMRI acquisition, the task-induced suppression of alpha power was inversely related to BOLD-activation in the contralateral sensorimotor cortex (Formaggio et al., 2010; Zich et al., 2015), highlighting that alpha-band oscillations are particularly sensitive to motor-related brain dynamics, making alpha power a reliable marker for motor function. However, while these earlier studies showed the utility of simultaneous EEG-fMRI, they were limited to analyzing EEG and fMRI separately and subsequently correlating them, without exploiting the full temporal dynamics and focusing on spatially restricted regions of interest rather than providing a whole-brain analysis that would allow a better comprehension of the broader neural networks involved. Only one work, to our knowledge, performed an EEG-driven fMRI analysis exploring BOLD correlates of EEG neural rhythms of a right-hand motor execution task (Sclocco et al., 2014). Our work aims to fill these gaps by applying a data-driven EEG-informed fMRI framework to motor tasks that include both execution and imagery of right and left hand. By using alpha power from bilateral motor regions, namely C3 and C4, we sought to identify the hemodynamic correlates of both ME and MI in an unbiased and comprehensive manner. This approach has the advantage of capturing the dynamic interplay between electrophysiological and hemodynamic signals across the entire brain without relying on predefined regions of interest. The primary aim of the present analysis was to cross-validate the EEG-informed fMRI results by comparison with conventional block fMRI analyses, which serve as the gold standard in detecting motor-related brain activity. Indeed, our findings indicated a predominant overlap between the EEG-driven fMRI maps and the fMRI activation maps in the sensorimotor regions like BA3, BA4, and BA6 during both execution and imagery tasks. These results align with the typical contralateral activation seen in motor tasks and reinforce the inverse relationship between alpha power suppression and BOLD signal increases during ME and MI tasks (Formaggio et al., 2010; Sclocco et al., 2014; Yuan et al., 2010; Zich et al., 2015). Besides sensorimotor regions, alpha power suppression was correlated with the cerebellum and insular cortex for motor execution and superior frontal gyrus for motor regions, regions further overlapped with fMRI results. In the MI condition, alpha-related activity was also seen in visual cortex areas, which likely reflects engagement of visual processing when imagining motor acts, as found in traditional fMRI analysis. More interestingly, these key motor regions overlapped, which gives stronger evidence that alpha-band oscillations are tightly coupled to sensorimotor cortex activities during motor tasks, which EEG-informed fMRI can successfully capture. Quantitative measures such as DC, precision, and recall also agreed on the consistency of findings between the two approaches. The precision of EEG-informed fMRI was high, indicating the regions identified by the EEG-driven model were highly specific to those activated in fMRI, although the sensitivity (recall) is a bit lower, reflecting that fMRI has detected some areas that are not captured by the EEG-informed approach, probably due to the regressor ability to capture only a part of the brain activity. These results confirm that EEG-informed fMRI represents a valid, block fMRI alternative but with the advantage of bearing electrophysiological information. Beyond the primary sensorimotor areas, our EEG-informed fMRI analysis revealed additional coactivated regions that were not shown in the fMRI analysis. For motor execution, this included the right angular gyrus (BA39), right temporal lobe (BA21, BA22), and right middle frontal gyrus (BA8, BA9) for right-hand movement, and the left inferior frontal gyrus (BA47) for left-hand execution. These may reflect additional parts of the network that also are involved in motor planning, integration of sensory feedback, and higher-order aspects of cognition associated with executing motor tasks (Farrer et al., 2008; Haggard, 2008; Hanakawa et al., 2008; Tankus & Fried, 2012).

The integration of EEG temporal dynamics with fMRI offers significant advantages in understanding the neural mechanisms underlying motor tasks. By focusing on alpha power suppression, we were able to map electrophysiological correlates of motor execution and imagery with high spatial specificity. Importantly, in the present study, our approach was data-driven and did not require any a priori assumptions about the tasks performed and the location of motor-related activations, demonstrating its potential to reveal novel patterns of brain activity. Our findings contribute to the growing body of evidence supporting the inverse relationship between alpha-band power and motor-related BOLD responses. Different activation patterns observed for ME and MI tasks suggest that while both engage overlapping sensorimotor networks, motor imagery may recruit additional cognitive processes, such as visual and associative areas, potentially reflecting the mental rehearsal of movement. Moreover, this illustrates the ability of EEG-informed fMRI to discriminate between right and left-hand tasks without explicit knowledge of sensor locations and task timings, highlighting its utility in multimodal neuroimaging studies.

The present findings have implications for sensorimotor rhythm brain-computer interface research using motor execution and imagery (Edelman et al., 2024; He et al., 2015; Yuan & He, 2014). In these BCI research, EEG rhythms during alpha band are often used to extract and decode brain intention to control a virtual or physical object. The present study provides neuroimaging basis on the high (negative) correlation of EEG alpha band power with BOLD responses in the sensorimotor area, which will aid in refining the design of BCI systems based on MI and ME paradigms. With the established EEG-informed fMRI mapping of the underlying brain activation using left and right-hand ME and MI, further investigation may explore more detailed somatotopic mapping of ME and MI as applied in more precise BCI control.

Despite the strength of our EEG-informed fMRI approach, several limitations need to be addressed. First, the dataset was not balanced for handedness. Although this issue was addressed in the unimodal fMRI analysis by including handedness as a covariate, it may still have influenced the motor imagery EEG-informed results. While our model successfully discriminated between right and left-hand motor execution, no significant results were found for left-hand motor imagery. Aside from the small dataset retained for analysis, the physiologically lower activation during left-hand motor tasks and the limited number of motor imagery trials in the protocol may have reduced the statistical power required for significant findings. To address these limitations, future studies should include a larger dataset and more trials. Lastly, we focused only on alpha power, which constrained our ability to investigate other oscillatory correlates involved in motor tasks.

## 5. Conclusion

In conclusion, this study demonstrates the utility of EEG-informed fMRI for investigating the neural correlates across multiple motor tasks without priori information on sensor locations or task timings, enabling temporally continuous analysis with full brain coverage. By integrating EEG alpha-band dynamics with BOLD signals, we were able to map motor-related brain activity with high spatial and temporal resolution. Our findings highlight the distinct yet overlapping brain networks involved in motor execution and imagery and underscore the potential of EEG-informed fMRI for advancing our understanding of motor functions and their potential applications, particularly in brain-computer interfaces and neurorehabilitation.

## CRediT authorship contribution statement

Elena Bondi: Writing – original draft, Visualization, Methodology, Investigation, Data Acquisition, Formal analysis, Conceptualization. Yidan Ding: Writing – original draft, Visualization, Methodology, Investigation, Data Acquisition, Formal analysis, Conceptualization. Yisha Zhang: Data Acquisition. Eleonora Maggioni: Writing – review & editing, Methodology. Bin He: Writing – review & editing, Methodology, Investigation, Supervision, Project administration, Conceptualization.

## Acknowledgment

This work was supported in part by NIH NS096761, NS127849, NS131069, and NS124564. The authors are grateful to Joe Disu and Dr. Sossena Wood for sharing equipment and useful discussions, and the CMU-Pitt BRIDGE Center for assistance in data acquisition. EM was supported by the Next Generation EU (PRIN 2022 PNRR, P20229MFRC).

## Data availability

The data supporting the conclusions are included in the paper and supplementary materials. Additional deidentified human data will be made available in a public online repository upon publication of the paper.

