## Supplementary Materials for "EEG-Informed fMRI Analysis Reveals Neurovascular Coupling in Motor Execution and Imagery"

### Supplementary Material

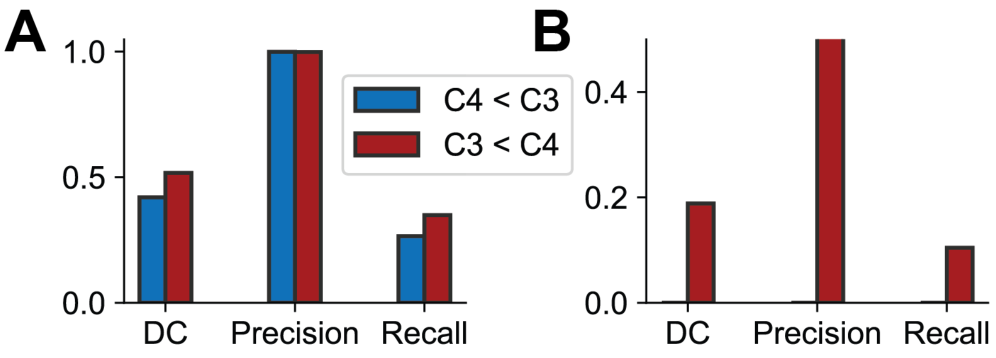


**Figure S1.**

**A:** Quantitative overlap between fMRI analysis and EEG-informed fMRI analysis results for ME tasks, constrained within the contralateral sensorimotor region (precentral and postcentral gyri), presented as dice coefficient (DC), precision, and recall. **B:** Quantitative overlap between fMRI analysis and EEG-informed fMRI analysis results for MI tasks, constrained within the contralateral sensorimotor region (precentral and postcentral gyri), presented as dice coefficient (DC), precision, and recall.

*Table S1. Group-level unimodal fMRI activations for motor execution (ME) using a random effect model with age, sex, and handedness as covariates (p < 0.001, k ≥ 15 voxels) (n = 17). Labeling performed with AAL3 atlas.*

**ME RH > ME LH**

| T-value (p<0.001, k > 15) | Cluster-size [# voxels] | Peak coordinates (xyz) [mm] | Atlas regions (%) |
|---|---|---|---|
| 11.0 | 981 | -40 -22 60 | Left Postcentral gyrus (63), Left Precentral gyrus (36) |
| 7.8 | 95 | -16 -22 4 | Left Ventral posterolateral (49), Left Pulvinar lateral (23), Left Intralaminar (11), Left Mediodorsal lateral parvocellular (7), Left Pulvinar medial (5), Left Ventral lateral (4) |
| 7.7 | 485 | 12 -50 -14 | Right Lobule IV-V of cerebellar hemisphere (80), Lobule IV-V of vermis (7), Right Lobule VI of cerebellar hemisphere (5), Right Lingual gyrus (4), Right Fusiform gyrus (3) |
| 7.3 | 87 | -32 -10 0 | Left Lenticular Nucleus-Putamen (68) |
| 6.8 | 54 | -38 -22 22 | Left Rolandic operculum (63), Left Insula (35), Left Postcentral gyrus (2) |
| 5.6 | 71 | -2 -16 54 | Left Supplementary motor area (89), Left Paracentral lobule (11) |
| 4.8 | 48 | 14 -62 -52 | Right Lobule VIII of cerebellar hemisphere (88), Right Lobule IX of cerebellar hemisphere (10) |

**ME LH > ME RH**

| T-value (p<0.001, k > 15) | Cluster-size [# voxels] | Peak coordinates (xyz) [mm] | Atlas regions (%) |
|---|---|---|---|
| 16.0 | 770 | -12 -48 -16 | Left Lobule IV-V of cerebellar hemisphere (71), Left Lobule VI of cerebellar hemisphere (18), Left Fusiform gyrus (6), Lobule IV-V of vermis (2), Left Lingual gyrus (2) |
| 12.0 | 669 | 30 -6 0 | Right Lenticular Nucleus-Putamen (32), Right Rolandic operculum (21), Right Insula (10), Right Lenticular Nucleus-Pallidum (7) |
| 12.0 | 1640 | 32 -24 64 | Right Postcentral gyrus-Putamen (70), Right Precentral gyrus (21), Right Superior parietal gyrus (4) |
| 9.0 | 112 | 8 -10 52 | Right Supplementary motor area (100) |
| 8.4 | 99 | 16 -22 4 | Right Pulvinar lateral - Ventral Posterior Medial Nucleus (26), Right Intralaminar (24), Right Ventral posterolateral (22), Right Pulvinar medial (10), Right Mediodorsal lateral parvocellular (9), Right Mediodorsal medial magnocellular (7), Right Ventral lateral (1) |
| 6.6 | 47 | -6 -66 -34 | Left Lobule VIII of cerebellar hemisphere (89), Left Crus II of cerebellar hemisphere (4) |
| 6.1 | 206 | -18 -58 -50 | Left Lobule VIII of cerebellar hemisphere (94), Left Lobule IX of cerebellar hemisphere (6) |
| 5.0 | 43 | 6 -22 50 | Right Supplementary motor area (86), Right Paracentral lobule (12), Right Middle cingulate & paracingulate gyri (2) |
| 4.7 | 28 | 24 -12 62 | Right Precentral gyrus (36), Right Superior frontal gyrus-dorsolateral (25) |

*Table S2. Group-level unimodal fMRI activations for motor imagination (MI) using a random effect model with age, sex, and handedness as covariates ($p < 0.001$, $k \geq 15$ voxels) ($n = 17$). Labeling performed with AAL3 atlas.*

**MI RH > MI LH**

| T-value (p<0.001, k > 15) | Cluster-size [# voxels] | Peak coordinates (xyz) [mm] | Atlas regions (%) |
|---|---|---|---|
| 8.3 | 19 | -2 12 -8 | Left Olfactory cortex (63), Left Nucleus accumbens (32), Right Olfactory cortex (3) |
| 6.3 | 29 | -46 -34 32 | Left SupraMarginal gyrus (34), Left Rolandic operculum (3) |
| 6.2 | 32 | -14 -84 14 | Left Cuneus (50), Left Calcarine fissure and surrounding cortex (41), Left Superior occipital gyrus (6) |
| 6.2 | 36 | -6 -40 32 | Left Middle cingulate & paracingulate gyri (86) |
| 5.8 | 50 | -26 -12 66 | Left Precentral gyrus (100) |
| 5.4 | 226 | -28 -28 60 | Left Postcentral gyrus (70), Left Precentral gyrus (18), Left Superior parietal gyrus (10) |
| 4.6 | 33 | 24 -64 -16 | Right Lobule VI of cerebellar hemisphere (64), Right Lingual gyrus (18), Right Fusiform gyrus (9), Right Lobule IV-V of cerebellar hemisphere (9) |

**MI LH > MI RH**

| T-value (p<0.001, k > 15) | Cluster-size [# voxels] | Peak coordinates (xyz) [mm] | Atlas regions (%) |
|---|---|---|---|
| 6.2 | 59 | 32 -6 -2 | Right Lenticular Nucleus-Putamen (88), Right Lenticular Nucleus-Pallidum (8) |
| 5.7 | 54 | 30 -22 58 | Right Postcentral gyrus (63), Right Precentral gyrus (35) |
| 4.9 | 24 | 38 -14 56 | Right Precentral gyrus (100) |

*Table S3. Group-level unimodal fMRI activations for motor execution (ME) using a fixed effects model (p < 0.001, k ≥ 15 voxels) (n = 8). Labeling performed with AAL3 atlas.*

**ME RH > ME LH**

| T-value (p<0.001, k > 15) | Cluster-size [# voxels] | Peak coordinates (xyz) [mm] | Atlas regions (%) |
|---|---|---|---|
| 34.0 | 3057 | -42 -22 62 | Left Postcentral gyrus (54), Left Precentral gyrus (34), Left Superior parietal gyrus (7), Left Paracentral lobule (1), Left Superior frontal gyrus-dorsolateral (1) |
| 14.0 | 1100 | 18 -48-22 | Right Lobule IV-V of cerebellar hemisphere (54), Right Lobule VI of cerebellar hemisphere (15), Lobule IV-V of vermis (13), Right Lingual gyrus (7), Right Fusiform gyrus (6), Lobule VI of vermis (4), Right Lobule III of cerebellar hemisphere (2) |
| 12.0 | 445 | 12 -62-46 | Right Lobule VIII of cerebellar hemisphere (79), Right Lobule IX of cerebellar hemisphere (15), Lobule VIII of vermis (4) |
| 7.8 | 364 | -2 -8 52 | Left Supplementary motor area (66), Left Paracentral lobule (28), Left Middle cingulate & paracingulate gyri (2) |
| 6.8 | 114 | -14 -22 6 | Left Ventral posterolateral (38), Left Pulvinar lateral (23), Left Intralaminar (15), Left Pulvinar medial (11), Left Mediodorsal lateral parvocellular (10), Left Ventral lateral (3), Left Mediodorsal medial magnocellular (2) |
| 6.4 | 97 | -30 -10 0 | Left Lenticular Nucleus-Putamen (62), Left Insula (2) |
| 5.6 | 148 | -36 -18 20 | Left Rolandic operculum (60), Left Insula (26), Left Postcentral gyrus (14) |

**ME LH > ME RH**

| T-value (p<0.001, k > 15) | Cluster-size [# voxels] | Peak coordinates (xyz) [mm] | Atlas regions (%) |
|---|---|---|---|
| 34.0 | 4108 | 38 -24 66 | Right Postcentral gyrus (42), Right Precentral gyrus (25), Right Supplementary motor area (10), Right Superior parietal gyrus (5), Right Superior frontal gyrus-dorsolateral (3), Right Paracentral lobule (2), Right Middle cingulate & paracingulate gyri (1) |
| 16.0 | 1846 | -16 -50 -20 | Left Lobule IV-V of cerebellar hemisphere (40), Left Lobule VIII of cerebellar hemisphere (23), Left Lobule VI of cerebellar hemisphere (16), Left Lingual gyrus (4), Left Fusiform gyrus (4), Left Lobule IX of cerebellar hemisphere (4), Lobule IV-V of vermis (3), Lobule VI of vermis (1), Lobule VIII of vermis (1) |
| 8.0 | 864 | 38 -16 20 | Right Rolandic operculum (33), Right Lenticular Nucleus-Putamen (27), Right Insula (14), Right Lenticular Nucleus-Pallidum (3), Right Heschls gyrus (1), Right SupraMarginal gyrus (1) |
| 6.5 | 91 | 16 -20 8 | Right Ventral posterolateral (27), Right Pulvinar lateral (26), Right Intralaminar (15), Right Pulvinar medial (14), Right Mediodorsal lateral parvocellular (9), Right Ventral lateral (3), Right Mediodorsal medial magnocellular (3), Right Pulvinar inferior (1) |

*Table S4. Group-level unimodal fMRI activations for motor imagination (MI) using a fixed effects model (p < 0.001, k ≥ 15 voxels) (n = 8). Labeling performed with AAL3 atlas.*

**MI RH > MI LH**

| T-value (p<0.001, k > 15) | Cluster-size [# voxels] | Peak coordinates (xyz) [mm] | Atlas regions (%) |
|---|---|---|---|
| 17.0 | 3547 | -38 -22 58 | Left Postcentral gyrus (48), Left Precentral gyrus (28), Left Superior parietal gyrus (11), Left Inferior parietal gyrus-excluding supramarginal and angular gyri (4), Left Paracentral lobule (3), Left Superior frontal gyrus-dorsolateral (2) |
| 7.2 | 571 | 16 -50 -20 | Right Lobule IV-V of cerebellar hemisphere (70), Right Lobule VI of cerebellar hemisphere (15), Right Fusiform gyrus (6), Lobule IV-V of vermis (4), Right Lingual gyrus (2) |
| 6.8 | 273 | 12 -60 -46 | Right Lobule VIII of cerebellar hemisphere (76), Right Lobule IX of cerebellar hemisphere (23) |
| 5.8 | 223 | -2 -10 52 | Left Supplementary motor area (83), Left Paracentral lobule (17) |
| 5.6 | 154 | -30 -6 -2 | Left Lenticular Nucleus-Putamen (59), Left Insula (9) |
| 5.0 | 245 | -46 -20 18 | Left Rolandic operculum (46), Left SupraMarginal gyrus (24), Left Postcentral gyrus (23), Left Insula (6) |
| 4.6 | 43 | 26 68 8 | Right Superior frontal gyrus-dorsolateral (93), Right Superior frontal gyrus-medial (7) |
| 4.5 | 32 | 6 8 -8 | Right Nucleus accumbens (84), Right Olfactory cortex (3) |
| 4.5 | 77 | 0 -38 28 | Left Middle cingulate & paracingulate gyri (49), Left Posterior cingulate gyrus (48), Right Middle cingulate & paracingulate gyri (3) |
| 4.3 | 285 | 40 40 38 | Right Superior frontal gyrus-dorsolateral (53), Right Middle frontal gyrus (47) |
| 3.7 | 45 | -14 -78 -18 | Left Lobule VI of cerebellar hemisphere (56), Left Lingual gyrus (22), Left Crus I of cerebellar hemisphere (22) |
| 3.6 | 27 | -16 -24 8 | Left Mediodorsal lateral parvocellular (44), Left Pulvinar lateral (30), Left Ventral posterolateral (19), Left Intralaminar (4), Left Pulvinar medial (4) |

**MI LH > MI RH**

| T-value (p<0.001, k > 15) | Cluster-size [# voxels] | Peak coordinates (xyz) [mm] | Atlas regions (%) |
|---|---|---|---|
| 17.0 | 1565 | 38 -20 58 | Right Postcentral gyrus (53), Right Precentral gyrus (36), Right Superior parietal gyrus (1) |
| 5.4 | 195 | -16 -48 -22 | Left Lobule IV-V of cerebellar hemisphere (91), Left Fusiform gyrus (7), Left Lingual gyrus (2) |
| 4.9 | 150 | 10 -8 52 | Right Supplementary motor area (100) |
| 3.8 | 25 | 30 -6 0 | Right Lenticular Nucleus-Putamen (100) |
